# Epigenetic Landscape Models: The Post-Genomic Era

**DOI:** 10.1101/004192

**Authors:** J Davila-Velderrain, JC Martinez-Garcia, ER Alvarez-Buylla

## Abstract

Complex networks of regulatory interactions orchestrate developmental processes in multicellular organisms. Such a complex internal structure intrinsically constrains cellular behavior allowing only a reduced set of attainable and observable cellular states or cell types. Thus, a multicellular system undergoes cell fate decisions in a robust manner in the course of its normal development. The epigenetic landscape (EL) model originally proposed by C.H. Waddington was an early attempt to integrate these processes in a universal conceptual model of development. Since then, a wealth of experimental data has accumulated, the general mechanisms of gene regulation have been uncovered, and the placement of specific molecular components within modular gene regulatory networks (GRN) has become a common practice. This has motivated the development of mathematical and computational models inspired by the EL aiming to integrate molecular data and gain a better understanding of development, and hopefully predict cell differentiation and reprogramming events. Both deterministic and stochastic dynamical models have been used to described cell state transitions. In this review, we describe recent EL models, emphasising that the construction of an explicit landscape from a GRN is not the only way to implement theoretical models consistent with the conceptual basis of the EL. Moreover, models based on the EL have been shown to be useful in the study of morphogenic processes and not just cell differentiation. Here we describe the distinct approaches, comparing their strengths and weaknesses and the kind of biological questions that they have been able to address. We also point to challenges ahead.

## Introduction

The development of multicellular organisms comprises cellular differentiation and morphogenesis (spatio-temporal arrangement of different cell types). The progressive loss of potency from pluripotent stem cells to mature, differentiated cells, as well as the reproducible emergence of spatiotemporal patterns through the course of development has been always perceived as strong evidence of the robustness and *deterministic* nature of development. The explanation of such a robust process has always puzzled researchers. For a long time, although not always stated explicitly, the prevailing paradigm in developmental biology was supported on two fundamental paradigms: (1) a mature cell, once established, displays an essentially irreversible phenotype; and (2) the developmental process is controlled by a “program” as a genomic blueprint following a simplistic linear scheme of causation in an essentially deterministic fashion. Discoveries in the last decade have challenged these assumptions. It has been shown that a differentiated state of a given cell is not as irreversible as previously thought, and that in fact it is possible to reprogram differentiated cells into pluripontent states with a plethora of protocols [1–3]. On the other hand, technological advances in time-lapse microscopy and live imaging allowed the *in-vivo* exploration of the role of physical constraints and non-genetic heterogeneity in tissue patterning and morphogenesis [4–6]. There is now compelling empirical evidence supporting intrinsic physical processes as a fundamental source of order (mainly ruled by stochastic self-organization processes) instead of deterministic pre-programmed rules [7]. Although these observations have just recently shift the focus of study of developmental biology and biomedical research, the new evidence supports the proposals that visionary theoretical biologists posited decades ago [8–12]. C.H. Waddington was one of the first to point out that the physical implementation of the information coded in the genes and their interactions imposes developmental constraints while forming an organism. The heuristic model of the epigenetic landscape (EL) was a visionary attempt to consolidate these ideas in a conceptual framework that allows to discuss in an intuitive manner the relationship between genetics, development, and evolution [8,13,14].

Now in the data-rich, post-genomic era the EL has been consolidated as the preferred conceptual model for the discussion of the mechanistic basis underlying cellular differentiation, trans-differentiation, and reprogramming events [15–17]. The holy grail of this prolific area of research, the so-called stem cells systems biology, is the possibility of discovering efficient reprogramming or therapeutic strategies by combining mathematical and computational modeling with experimental techniques [18–22]. The formalization of the EL in the context of the study of the dynamical properties of GRNs is a salient theoretical framework which is starting now to show great promise in this direction. Given the current relevance of the intended purpose of modeling the EL associated to molecular networks involved on processes such as stem cell differentiation [23], tissue morphogenesis [24], and carcinogenesis [25]; and the fact that different approaches have been proposed in order to reach similar goals [26–28], here we present an integrative and comparative review which considers the most successful approaches used up to now. Our objective is two-fold: (1) to help different research groups attempting to formalize the EL to reduce the gap existing between current different approaches and (2) to contribute to shape a common discussion ground on EL models between experimentalists and theorists. With this in mind, we present both a conceptual and rigorous exposition of otherwise rather technical and disjunct subjects.

In the sections that follow, we first explain briefly the dynamical systems view of the cell, *i.e*. the standard theoretical framework adopted for the mathematical and computational study of cell differentiation. This formal conceptualization of the cell gives rise to the rigorous definition of a given cell phenotype and is the starting point for the construction of EL models. We also elaborate on why there is a growing interest in extending GRN models towards the implementation of EL models. We then summarize some recent work in which derivations of the EL associated with gene regulatory circuits and networks are presented. Moreover, we discuss both the biological and mathematical difficulties associated with each of the considered approaches that have been used. Finally, we conclude with some perspectives and open problems regarding the generalization of computational models motivated by the conceptual basis of EL models.

## The Dynamical-Systems View of Cell Biology

Recently, a modern picture of the EL has been emerging in the context of GRNs [27]. An assumption of the “systems dynamics” approach to biology is that cell behavior can be understood in terms of the dynamical properties of the involved molecular regulatory networks [29,30]. This idea, particularly in the context of development, aligns very well with Waddington’s intuition [15], and is now supported by a wealth of consolidated theoretical and experimental work [31,32]. As a consequence, the orchestrating role of GRNs in developmental dynamics is now widely accepted [33–36]. Furthermore, with the use of modern experimental techniques, the number of known specific gene interactions is growing, and the mechanisms of gene regulation are fairly well understood. This knowledge enables the mathematical description of experimentally grounded GRNs in terms of dynamical systems. There are several excellent reviews on the dynamical-systems view of cell biology [24,33,37–39]; in this section, we briefly explain this view.

Under dynamical-systems framework, a cell is considered a dynamical system, assuming that its state at a certain time can be described by a set of time-dependent variables. As a first approximation, it is assumed that the amount of the different proteins or the levels of gene expression (e.g. expression profiles) are sufficient to describe the cellular phenotype [40]. Thus, the expression profile is taken as the set of variables representing the state of the cell; each gene in the cell’s GRN represents one variable (see Figure 1). Mathematically, the set of variables is represented as a state vector given by x(*t*)=[*x*_1_(*t*), *x*_2_(*t*)…, *x*_*n*_(*t*)] for a GRN with *n* genes; conceptually, this vector represents the corresponding phenotype. It is useful, both conceptually and mathematically, to imagine an abstract space termed the *state space* of the system, which in the context of GRNs corresponds to the abstract space comprising all the theoretically possible phenotypes a cell can exhibit; each point in this abstract space represents one particular phenotype of the cell that specifically corresponds to a configuration or vector with the expression states or levels in discrete and continuous models, respectively, for each molecular component (node) considered in the network (Figure 1b). Furthermore, it is assumed that the phenotype at a certain time and the phenotype at a later time are connected by a state trajectory in a causal way. Mathematically, the current phenotype is a function or a more general mapping of the initial phenotype and certain additional parameters, which may take into account the influence of the environment and possibly also actions of external control. The connection between phenotypes can be formally expressed by a dynamical equation,

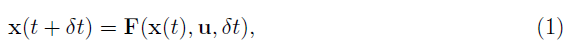

**Figure 1.**
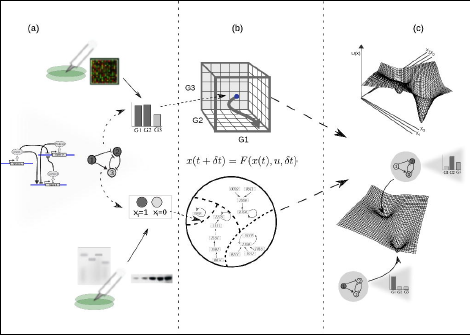
**From experimental data on gene function and interactions to a dynamic gene regulatory network and epigenetic landscape model.** (a) The architecture of a GRN is proposed given available experimental molecular data; the state of the network is specified as a gene expression profile or gene on/off (1/0) configuration for the case of continuous or discrete state models, respectively. Boolean or differential equations are used, respectively. (b) The complete set of possible states define a continuous (above) or discrete (below) state space, where each state corresponds to a point; changes in gene expression during developmental dynamics manifest as trajectories in this abstract space (here depicted as arrows). (c) In an intuitive characterization of an EL, an ‘elevation” value *U*(*x*) is associated to each network state *x*. The association of ‘elevation” values to network states, or more generally, the quantitative characterization of their relative stability is the ultimate goal of EL modeling efforts. For illustrative purposes, the EL is depicted here as a hypothetical low-dimensional projection.

where **F** represents the map that connects one phenotype with immediately previous phenotype (**F** is also known as the *transition map),* x(*t*) denotes the phenotype at a certain time *t*, and **u** stands for the vector of additional parameters. Both the time increments *δt* and the state variables *x*_*i*_(*t*) can be either continuous or discrete, depending on the chosen mathematical formalism used. Within the cell, the map **F** is implemented by the architecture of the GRN, which specifies both the topology of the network and the nature and form of the corresponding gene regulations [26]. Thus, through the causal connections between the phenotypes, the GRN imposes dynamical constraints and limits the permissible behavior of the cell, specifically of the transient and stable configurations that it may attain. The existence of a dynamical map, which defines the phenotype at a certain time, by means of the corresponding phenotype at an earlier time, expresses the causality of the cellular developmental process. We must point out that when modeling developmental dynamics, two different cases of causal relationships can be considered between two given phenotypes. If the relationship between the two given phenotypes is unambiguous, *e.g.* if the future phenotype is given by the map in a unique way by the initial one, the corresponding model is deterministic. In contrast, if there are several possibilities for the future phenotype depending on *random effects,* the model is stochastic. One of the most salient and impressive features of GRNs is the existence of a small number of stationary gene configurations within the state space [10]. Given a specific GRN, a specific set of phenotypes could satisfy the constraints imposed by the GRN, that is, each of these phenotypes would be connected to itself by the map **F**, *e.g.* **x**\*=**F**(**x**\*, **u**). When these steady states (**x**\*) are also resilient to perturbations, that is, if they return back to the steady state after being kicked away by state variations either of intrinsic or external origin, we refer to them as *attractors*; and the theory states that these correspond to the observable phenotypes or cell types [33]. In formal terms, an attractor state is both stationary (time-invariant) and stable [40]. All other states are either unstable or form part of transitory trajectories channeled towards one of these attractor states. From a geometric point of view, attractors represent regions in the state space toward which the system trajectories are channeled to due to dynamic constraints imposed by the GRN (see [41] for a through and intuitive explanation).

In the last two decades, researchers in the area of systems biology have witnessed the great success of GRN dynamical models as integrative and explanatory frameworks for specific developmental processes. Available experimental techniques in molecular biology enable the physical characterization of regulatory interactions, and from these it is possible to postulate experimentally grounded GRN models which translate to mapping functions **F** in a dynamical model. The study of the resulting GRNs of specific real developmental processes has shown that the emergent attractor states of the models are indeed consistent with the observable cell type phenotypes (expression configurations or profiles) in both wild-type and mutant backgrounds (see, for example: [35,36,42–44]). Experimental evidence supporting the hypothesis of cell fates as high-dimensional attractors has also been presented [31,32]. The role of experimentally controllable parameters in differentiation and cell-fate decision events is also commonly studied [45–48]. Thus, it seems that the postulation of experimentally grounded GRN dynamical models, their qualitative analysis and dynamical characterization in terms of control parameters, and the validation of predicted attractors against experimental observations has become a well-established, successful framework for the study of developmental dynamics [49] (Figure 1a–b).

### Extending GRN to EL Models

The qualitative analysis of the dynamics of GRN models is perfectly suited for the study of the specification of cell fates as a result of the constrains imposed by the associated GRN. This conventional analysis includes the identification and local characterization of attractor states and the comparison of these predicted cell types configurations with the ones that are actually observed in the corresponding biological system. If one is interested in studying the potential transition events among the already characterized stable cellular phenotypes, however, several difficulties arise. Standard analysis of dynamical systems, which focuses on the existence and local properties of a given attractor, fail to capture the main problem which is concerned with the relative properties of the different attractors [28]. In deterministic GRN models, given certain values for the related control parameters, the system under study always converges to a single attractor if initialized from the same state, and once it attains such steady-state it remains there indefinitely. In contrast, during a developmental process, cells change from one stable cell configuration to another one in specific temporal and spatial or morphogenic patterns. Thus, we need additional formalisms in order to explore questions regarding how cells in the course of differentiation decide between one of the available given attractors; or the order in which the system converges to the different attractors, given an initial condition; and to predict how these mechanisms can be altered by external control.

### Modeling Goals

The need for extending GRN dynamical models beyond standard local analysis is related with the interest in addressing the following - and similar - questions. Conceptually, given an experimentally determined GRN, how can we study and predict both specific “normal” and altered cellular differentiation events or morphogenic patterns? Is it possible to control the fate of differentiation events through well-defined stimuli? Can we deliberately cause altered morphogenic patterns by means of either genetic or environmental perturbations? Or formally, given a specific dynamical mapping **x**(*t*+*δt*)=**F**(**x**(*t*), **u**, *δt*), and its associated state space, how can we study the conditions under which a transition event occurs among the attractor states **x**\*? Is there a reproducible pattern of transitions? Are we able to alter the observed pattern through specific external control perturbations **u**? To what extent are the observed temporal or spatial morphogenetic patterns emergent consequences of the GRNs? The extension of GRN dynamical models and their analysis in order to address these and similar questions has shown to be a fruitful area of research in recent years [23,25,28,50–54]. The conceptual basis for most of these efforts is the EL.

In the following sections we describe some of the different approaches that have been proposed recently in order to tackle the formulated questions. We start from the simplest deterministic approaches, which primary goal is to derive an explicit landscape (potential) function from the associated gene regulatory circuits. Then we focus on probabilistic approaches: these have shown to be the most appropriate framework to extend GRN dynamical models. Within the stochastic approaches, we distinguish those which attempt to derive an explicit probabilistic landscape from those that are concerned with the transition probabilities between attractors and the temporal evolution of the probability distribution over the cellular states. We give special attention to the kind of biological questions that may be better addressed by the different approaches reviewed here.

## Deterministic EL Models From Genetic Circuits

### An Introductory Toy Model

A quite simple auto-activating single-gene circuit, a basic model of cell differentiation induction, is exposed in [55] as a conceptual tool to discuss some difficulties regarding Waddington’s EL. In this work, the EL is mathematically described by a potential function. In dynamical systems theory, besides the state space approach explained briefly above, there is another way to visualize the dynamics of a system, but applicable only if the system is simple enough: the potential function [56,57]. The potential is a function *V(x)* which (in one-dimensional systems) fulfills the relation given by:

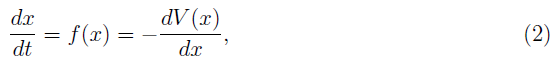

*i.e. f* (*x*) is the negative derivative of the potential, which can be found by direct integration:

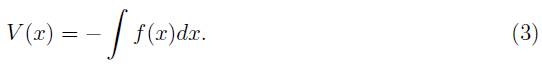

Such a function defines an attractor landscape for the given dynamical system, and its plot graphically represents the dynamics of the system (Figure 2). Specifically, *minima* of the potential correspond to fixed-point attractors (*e.g*. cell types), and *maxima* correspond to unstable fixed-points. The motion, *i.e.* the state trajectories are given by the gradient lines (the lines of steepest descent of the potential). The trajectories are attracted by the minima of the potential. This corresponds to an intuitive, direct derivation of the EL: a “height” value is associated to each of the points in the state-space in a way that those regions corresponding to attractors (cell types) will have a lower value than that of the other transitory states (Figure 2c). Conceptually, the rolling ball of Waddington’s EL will represent the state of a differentiating cell moving from higher to lower regions in state space. Thus, the calculated heights of the different attractors are expected to reflect their developmental potential in a hierarchical way: the lower height the lower potential for differentiation. All one-dimensional systems have a potential function, but most two- or higher-dimensional systems do not [57]. This means that one could only apply this method if the cell is represented by a single-gene (single variable) circuit. Furthermore, note that here the EL plays the role of a “toy” model useful in conceptual discussions, a role quite relevant (see [55]) but similar to that of the original metaphorical proposal of Waddington. In this review we devote more attention to the application of EL models to real specific developmental processes with explanatory and predictive purposes. A more “realistic” sub-network model incorporating several transcription factors in a modular structure would probably be necessary in such cases. The application of the integration-based potential function approach, however, cannot be applied to cases with a higher number of genes. Also one should be cautious when postulating the existence of a potential for living systems in strict sense: a cell is an open non-equilibrium thermodynamical system, and its dynamics in general does not follow a gradient (since the transition rate between two given attractor states is not path-independent) [28,40]. For this reason authors use the term “quasi-potential” when speaking about cellular dynamics from a system-dynamics point of view (see below). In many cases, in continuous-time models of GRNs the dynamics is given by more general types of autonomous differential equations (DEs) that cannot be derived from a potential function. The time evolution of the cell state x*(t)* is commonly modeled by the system of DEs:

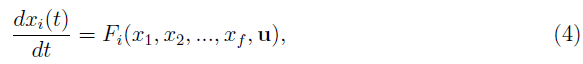

where *i*=1, 2,…, *n* for a GRN of *n* genes. A dynamics defined by such a general DE is a special form of the map in equation (1).

**Figure 2.**
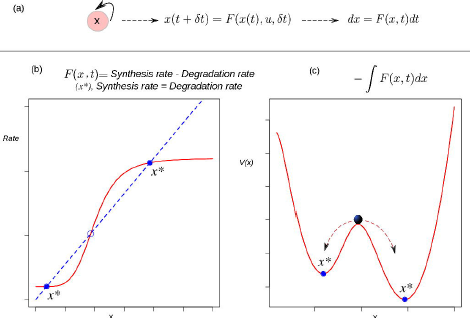
**The derivation of a potential function to visualize the epigenetic landscape and the dynamics of a system.** (a) The causal connection between the state of the system (an auto-activating gene circuit) at a certain time and its state at a later time is modeled by a differential equation. (b) Attractors in rate-balance analysis. The blue dotted line and the red curve represent respectively a linear degradation rate, and a nonlinear synthesis rate for the circuit’s gene; the restrictions imposed by the circuit to the systems dynamics are met when both rates are balanced. The states that meet this balanced condition are stationary, and if stable (filled circles), are denoted as attractor states *x*^*^. (c) The potential function. The attractor states *x*^*^ lie at the bottoms of valleys (minima). The trajectories starting from unstable, transitory states are attracted by the minima of the potential. The relative stability of the left (right) attractor with respect to the other is lower (higher) as quantified by the lower (higher) barrier height between them.

### A Deterministic Model of Generic Cell Fate Specification

In the past few years, it has been proposed that small gene circuits with a specific topology drive cell-fate decision at branch points of cell development [22]. One of these circuits, which has been extensively studied both experimentally and theoretically, is that in which the component genes have both cross-inhibitory and self-activating interactions [45,58,59]. Bhattacharya and collaborators recently proposed a numerical approach to derive a deterministic path-integral “quasi-potential” from this two-gene regulatory circuit as a general model of cell fate specification [60]. As mentioned in the previous section, the number of cases where it is possible to derive an analytical potential function is in fact very reduced: in general, the functions **F** in the continuous-time model for cellular dynamics (equation (4)) are nonlinear, and cannot be analytically integrated. In order to derive a function with similar analytical properties than the potential function, the authors proposed a computational-based strategy. They defined a quantity *U_q_* that changes incrementally along a trajectory in state space. According to the definition of a potential function (equation (3)), the change of such a quantity Δ*U_q_* for positive time increments is always negative along a trajectory. This ensures that trajectories flow downhill, and that the attractors of the system would correspond to local minima where the change in Δ*U_q_*=0. They termed the quantity (*U_q_*) as “quasi-potential” , and calculated its overall change by integrating numerically the change (Δ*U_q_*) along trajectories originating from different points in the phase space. The quasi-potential surface is then drawn graphically by interpolation of trajectories (see [60] for details). Among the principal contributions of this work, according to the authors, are the applicability of their approach to systems for which an analytical potential function cannot be derived as well as to high-dimensional GRNs. Although simple and intuitive, this computational approach suffers from several important limitations. The proposed quasi-potential is minimized along a trajectory *by construction.* It is not clear what information regarding the system can be obtained from this mapping which is not already attainable through the characterization of the 2-D phase space using standard analysis techniques from the qualitative theory of nonlinear dynamics [56,57]. On the other hand, if one considers a gene regulatory module whose typical number of composing genes range from 10 to 20 - but may contain more than 50 genes (see below) - even if theoretically possible, it is impractical to apply to these higher-dimensional systems either the proposed numerical integration approach or the proposed methodology for drawing the quasi-potential surface. For these and related reasons, the proposed deterministic “quasi-potential” has been criticized; for example, Zhou and collaborators noted that it does not represent either Lyapunov stability or transition rates among different attractors [28]. Consequently, even when a graphical EL is effectively drawn, the general questions raised in the previous section cannot be addressed using this approach.

## Stochastic EL Models From Boolean GRNs

The first computational model envisioned for the simulation and analysis of the dynamic behavior of GRNs was the Boolean Network (BN) model [10,11]. Such approach has been extended to model various developmental processes in the context of the EL [25,50,51, 61,62]. A BN models a dynamical system assuming both discrete time and discrete state. This is expressed formally with the mapping:

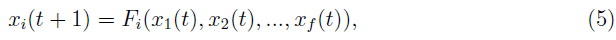

where the set of functions *F_i_* are logical prepositions expressing the relationship between the genes that share regulatory interactions with the gene *i*, and where the state variables *X_i_(t)* can take the discrete values 1 or 0 indicating whether the gene *i* is expressed or not at a certain time *t*, respectively. An experimentally grounded Boolean GRN model is then completely specified by the set of genes proposed to be involved in the process of interest and the associated set of logical functions derived from experimental data [24]. A dynamics defined by such a mapping is a special form of the map in equation (1).

### Attractor Transition Probability Approach to Explore The EL

In a deterministic framework once a cell state corresponds to an attractor, it will remain there indefinitely. The set of conditions that lead to each attractor comprise the attracting *basin.* Under stochastic fluctuations, the borders of attractor regions in state space may be reached and may be crossed, leading to transitions between attractors [63]. Thus, the implementation of an stochastic dynamical model opens the opportunity to study signal-independent transitions among attractors. There are several approaches to include probabilities in dynamical models. One approach is based on the idea of introducing transition probabilities. As discussed above, when studying cellular developmental dynamics, the transitions of interest are those among attractor states. Can these transitions be studied in terms of probabilities? The answer to this question is positive, since Boolean GRN can be extended in order to include stochasticity and then transition probabilities among attractors can be estimated. Several ways have been proposed to include stochasticity in a Boolean GRN model [64]. One way is the so-called stochasticity in nodes (SIN) model. Here, a constant probability of error *ξ* is introduced for the deterministic Boolean functions. In other words, at each time step, each gene “disobeys” its Boolean function with probability *ξ*. Formally:

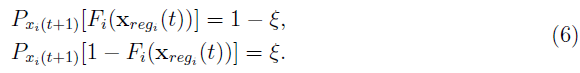

The probability that the value of the now random variable *x*_*i*_(*t*+1) is determined or not by its associated logical function *F*_*i*_(*x*_*reg*_*i*__(*t*)) is 1 — *ξ* or *ξ*, respectively.

Alvarez-Buylla and collaborators used this extended BN model to explore the EL associated with an experimentally grounded GRN [50]. In a BN model the set of possible states (or phenotypes) is finite. Specifically, the state space of a Boolean GRN with *n* genes has a size of 2*^n^* and is composed by the set of all possible binary vectors of length *n* (see Figure 3a). Thus, by simulating a stochastic one-step transition starting from each of the possible states a large number of times it is possible to estimate the probability of transition from an attractor *i* to an attractor *j* as the frequency of times the states belonging to the basin of the attractor *i* were mapped into a state within the basin of the attractor *j*. In [50], the authors followed this simulation approach to estimate a transition probability matrix ∏ with components:

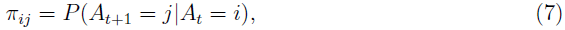

representing the probability that an attractor *j* is reached from an attractor *i* (Figure 3b). Once the set of attractors is known and the transition probability matrix is estimated, it is straightforward to implement a discrete time Markov chain model (DTMC) and obtain a dynamic equation for the probability distribution [65]. Indeed:

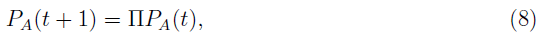

where *P*_*A*_(*t*) is the probability distribution over the attractors at time *t*, and ∏ is the transition probability matrix previously estimated. This equation can be iterated to simulate the temporal evolution of the probability distribution over the attractors starting from a biologically meaningful initial distribution. The extension of a Boolean GRN in order to apply this approach is quite simple and intuitive; however, there is a limitation that impedes its general applicability: as the size of the GRN grows, it is not possible to exhaustively characterize the attractor’s landscape associated with the GRN in terms of the emergent attractors and its corresponding basins of attraction. If the dynamics of the Boolean GRN is not exhaustively characterized, the corresponding transition probabilities among attractors cannot be estimated using the proposed approach.

**Figure 3.**
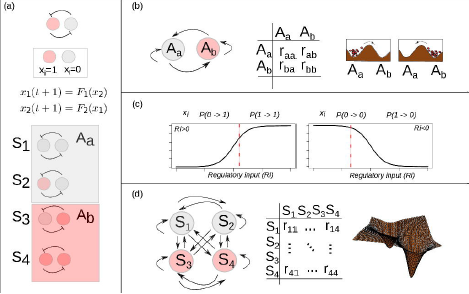
**Stochastic epigenetic landscape models from Boolean dynamics.** (a) A simple mutual-inhibition circuit is modeled as a Boolean network: discrete temporal evolution and binary (0,1) state variable. The discrete state space corresponds to the set of binary vectors (here 4 possible states) and is partitioned by two basins of attraction. (b) There are 4(2^2^) possible transitions among the two attractors. A 2 × 2 transition probability matrix specifies the probability of each possible transition. (c) Han and Wang proposed the use of a sigmoidal function of the total regulatory input (Ri) to calculate the probability of a one-step state transition of one gene *i* [51]. A specific value of the function (vertical red dotted line) gives the probability of the gene *i* becoming active (left) or inactive (right), given its regulatory input (Ri) in the current time. (d) There are 16(2^2^ × 2^2^) possible transitions among the 4(2^2^) possible states. A 2^2^ × 2^2^ transition probability matrix specifies the probability of each possible transition.

### EL Exploration in Flower Morphogenesis

Alvarez-Buylla and collaborators applied the previous approach to study the role of stochastic perturbations of the GRN underlying floral organ primordial cell fates in the observed robust temporal morphogenetic pattern observed in most flowering plant species [50]. In flowering plants, a floral meristem is sequentially partitioned into four regions from which the floral organ primordia are formed and eventually give rise to sepals in the outermost whorl, then to petals in the second whorl, stamens in the third, and carpels in the fourth whorl in the central part of the flower. This spatiotemporal pattern is widely conserved among angiosperms. Can the temporal pattern of cell-fate attainment be explained by the interplay of stochastic perturbations and the constraints imposed by a nonlinear GRN? Starting from the previously characterized Boolean GRN of organ identity genes in the *A. thaliana* flower [35], and applying the stochastic approach described above, the authors showed that the most probable order in which the attractors are attained, is in fact, consistent with the temporal sequence in which the specification of corresponding cellular phenotypes are observed *in-vivo*. The model provided then a novel explanation for the emergence and robustness of the ubiquitous temporal pattern of floral organ specification, and also allowed predictions on the population dynamics of cells with different genetic configurations during development [50]. Note that this approach constitutes a landscape-independent implementation of an EL model which successfully addressed an specific biological question; it also constitutes a new approach to understanding a morphogenic process. As seen in this example, it is important to stress that, through the calculation of transition probabilities among attractors, it is possible to explore the EL associated with a GRN independently of an explicit derivation of a landscape or quasi-potential. In the same paper, the authors also showed that a stochastic continuous approximation of the GRN under analysis yielded consistent results.

### Probabilistic Landscape (quasi-potential) Approach

Han and Wang proposed a different approach in order to extend a BN model. Their goal was to first estimate the one-step transition probabilities among all the possible points in the state space and not just among given attractors [51]. For this, they implemented a variation of the BN that was previously proposed by Li and collaborators [66] and which has been called the threshold network formalism [67]. In this model, the structure of the network is formally represented with an adjacency matrix C, whose components *c_ij_* take a value from the set [−1, 0,1] indicating the nature of the interaction - inhibition (−1), absence (0), or activation (1) - from the gene *j* to gene *i*. The dynamic mapping for the BN takes the form:

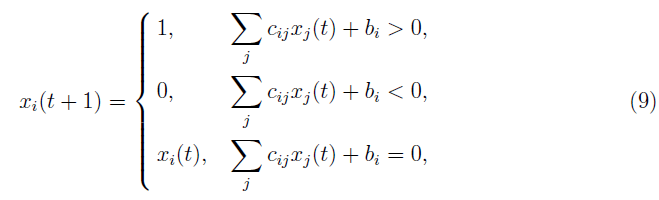

where *b_i_* is a parameter representing the ground state of the gene *i*: its state in the absence of regulation. In words, if the total input of a gene in the network is positive (activation), negative (repression) or zero; the future state of the gene will be active, inactive or unchanged from its previous state. Here, the total input of a gene is the sum of the previous states of the genes regulating it. The characterization of the entire attractor’s landscape can then be done through numerical iteration of this dynamical map as long as the network has a moderate size.Han and Wang extended this deterministic BN model into a probabilistic framework as follows. If the interest is focused on the computation of the probability of transition from one phenotype (state) to another phenotype for each of the 2*^n^* possible phenotypes in state space, then it is necessary to introduce a transition probability matrix with the probability of all possible transitions; this calculation is practically impossible. In order to work this limitation, Han and Wang introduced a simplification: they assumed that the one-step transition probability of one phenotype to another can be expressed as the product of the probability of each gene in the network being activated or not given the phenotype of the network in the previous time. Formally:

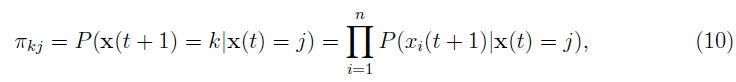

where *j* and *k* represent two different phenotypes and can take values from [1,…, 2*^n^*]; *n* is the number of genes in the network. The factorized transition probabilities are calculated by inserting a nonzero regulatory input 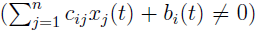 as the argument of a sigmoidal function whose range spans from 0 to 1, which is to say:

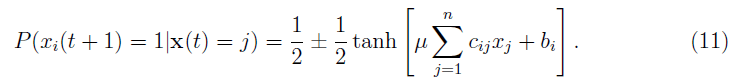

In the case of no input (*i.e.* 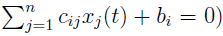 a small-valued parameter *d* is introduced:

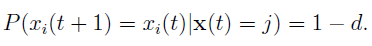

In words, the probability that a gene *i* will be on (1) at the future time *t*+1 will be closer to one as long as its total input at the previous time *t* is high. Similarly, the probability of being off (0) at the future time will be closer to 1 as long as the regulatory input is low (see Figure 3c). On the other hand, if there is no input to the gene, the probability of no change from its previous state is close to 1, and the closeness depends on the parameter *d*, a small number representing self-degradation. These rules ensure that the state of a gene will flip only if its total input is large enough.

After having calculated these probabilities, the general idea is then to use this information to obtain an appropriate “height” measure for each of the 2*^n^* states. With this in mind, the interest is first in calculating a steady-state probability distribution *P_SS_*(x). This stationary probability distribution is analogous to stationary configurations in the deterministic case; however, in the stochastic framework what keeps invariant in time is not the state of the system but the probability of being in any particular state. In other words, when this stationary distribution is reached, the probability of observing a cell in a particular phenotype does not change. Intuitively, one would expect that attractors would have a higher probability of being reached than transitory states. Thus, from a landscape perspective, the potency of differentiation and height should be inversely related with the probability. The approach that has been followed is to associate this *P_SS_*(x) with a height value. Wang has proposed that the probability distribution for a particular state *P*(x*_i_*)=exp[*U*(x*_j_*)], and from this expression then *U*(x*_i_*)=− ln *P*(x*_i_*), where *i*=1,…, 2*^n^*. This function *U* has been termed the (probabilistic) quasi-potential [26,54,68]. The association between such a “quasi-potential” and the steady-state probability is still an open research area (see below) [28]. The key point which has been championed by Wang and coworkers is that, although there is (in general) no potential function directly obtainable from the deterministic equations for a network, a generalized potential (or “quasipotential”) function can be constructed from the probabilistic description. This generalized potential function is inversely related to the steady-state probability [51,69,70]. For the case of the extended BN model, once the transition matrix is calculated, the information of the steady-state probabilities can be obtained by solving a discrete set of master equations (ME) for the network [51]. The so-called ME is a dynamical equation for the temporal evolution of a probability distribution [71]. In discrete form it is written:

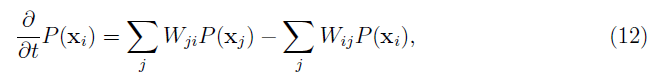

where we used *W_ij_* to denote the transition probabilities resulting from (11). The difference between this dynamical equation and the one discussed in the previous section is that here the time variable is treated as a continuous one. It is in general quite complicated to analyze MEs. In the case of this model, one ME is obtained for each of the 2*^n^* states. Han and Wang conveniently analyzed the whole set of equations following a numerical (iterative) method starting from uniform initial conditions *P*_x*i*_(*t*_0_)=1/2*^n^* and iterating until a stationary distribution is reached [51].

### From Probabilistic Landscapes to Putative Cancer Therapies

The previous approach has been applied to two specific processes: cell cycle regulation [51], and DNA damage response [25]. In the former case, the focus was on the global robustness properties of the network. Here we discuss the biological implications derived from the latter case. Choi and collaborators applied this BN probabilistic landscape approach to study state transition in a simplified network of the p53 tumor suppressor protein. The analysis of this network using the probabilistic landscape approach allowed the systematic search for combinatorial therapeutic treatments in cancer [72]. Given the network, key nodes and interactions that control p53 dynamics and the cellular response to DNA damage were identified by conducting single node and link mutation simulations; as a result, one network component, the molecule Wip1, was identified as one of the critical nodes. The flexibility of the BN model also enabled the specification of a MCF7 cancer cell by fixing the state of three nodes of the “normal” network in the course of simulations (for details, see [25,72]). Having specified two different network models, it was possible to compare the dynamics and associated quasi-potential of both normal cells and cancer cells in the absence and presence of DNA damage. Previous experimental observations indicated that prolonged p53 activity induces senescence or cell death; this behavior was shown to result from the inhibition of the interaction between the molecules Mdm2 and p53 caused by the action of the small molecule Nutlin-3 [73]. Using the model, Choi and collaborators predicted that neither Wip1 nor Mdm2-p53 interaction mutation alone were sufficient to induce cell death for MCF7 cancer cells in the presence of DNA damage; furthermore, the model provided a mechanistic explanation for this behavior: the effect of each of this perturbations alone is not enough to move the system out of an specific attractor’s basin. However, the simultaneous application of the two perturbations drives cancer cells to cell death or cell senescence attractors. These theoretical predictions were then validated using single-cell imaging experiments [25,72]. This study illustrated in an elegant way how cancer therapeutic strategies can be studied in mechanistic terms using a computational EL model. It must be pointed out that this result opened the door to the rational design of system dynamics cancer therapeutical techniques, in contrast to trial and error and reductionist approaches that have dominated the biomedical field.

## Stochastic EL Models From Continuous GRNs

Like the BN model, the general system of DEs can be extended in order to include stochasticity (in order to approach the modeling formalism to actual biological circunstances). The most intuitive extension considers the introduction of driving stochastic forces; in this approach, equation (4) is extended as:

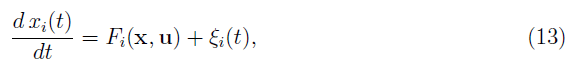

where ξ_*i*_(*t*) is the *i*th component of a driving stochastic force with zero mean value (*i.e.* < ξ_*i*_(*t*) >>= 0). This description, the so-called Langevin equation, is frequently used to model cellular dynamics under stochastic fluctuations [23,53,54,74]. Although intuitively simple at first sight, the consideration of a randomly varying quantity affecting the dynamics of the system implies several conceptual issues that should be considered with care. Any single cell will follow an erratic trajectory in state space, and its developmental dynamics will make each realization different *even if it starts from exactly the same initial condition.* Under this stochastic scenario, two equivalent perspectives to study the stochastic dynamics can be considered. On the one hand, the analysis could be focused on trajectories described by Langevin type equations, which describe the developmental dynamics of a single cell (Figure 4a). On the other hand, as the stochastic forces ξ_*i*_(*t*) vary from cell to cell in an *ensemble* (population) of cells, the state x(*t*) will also vary from cell to cell at any given time. One therefore may ask for the probability *P*(x, *t)* to find the state of a cell in a given state interval of the state space or, equivalently, for the frequency of cells in the ensemble whose states are in that state interval. The focus then shifts from the dynamics of the state of one cell to the dynamics of the distribution over the states in a given ensemble of cells. Indeed, an equation for the temporal evolution of this distribution *P*(x, *t*) can be constructed, the so-called *Fokker-Plank equation (FPE)* [75]:

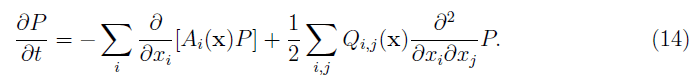

In mathematical terms, the corresponding process is known as a *diffusion process,* a mathematical model for stochastic phenomena evolving in continuous time [76,77]; the vector **A**(x) is known as the drift vector and the matrix **Q**(x) as the diffusion matrix. The FPE describes the change of the probability distribution of a cell state during the course of time (Figure 4b). Conceptually, the latter modeling perspective can be interpreted as the temporal evolution of a cloud (ensemble) of cells diffusing across the state space following both attracting and stochastic forces [78]. The stochastic nature of the trajectories also allows qualitatively richer dynamics in state space. Specifically, if one is interested in the developmental connection between one specific initial phenotype and one specific final phenotype - for example two different given attractors - there is no longer a single possible path connecting them. Instead, the same final phenotype can be reached following different paths in state space (Figure 4c). This situation raises yet another interesting problem: are all the paths equally probable? Is there a dominant path for the transition? Physicists have proposed the so-called path-integral formalism in order to tackle these and similar questions [79]. Specifically, one may want to answer what is the probability of starting from an initial phenotype at a certain time and ending in another phenotype at a future time. The conceptual basis of this strategy is based on the idea of calculating an average trajectory (*e.g.* integrating over the possible paths). The calculated averaged path corresponds to the dominant path that the underlying process follows (Figure 4c).

**Figure 4.**
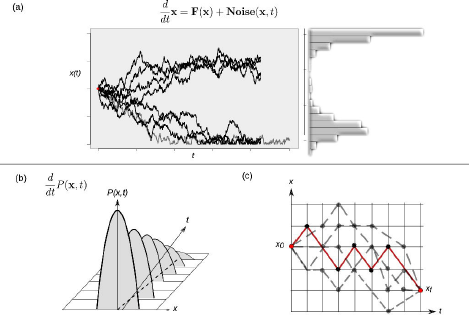
**Differente approaches to study continuous-time stochastic models of the epigenetic landscape and developmental dynamics.** (a) A continuous-time stochastic (diffusion) model is driven by a drift (deterministic) component **F** and an stochastic (**Noise**) force. The graph shows 10 different realizations of the stochastic dynamics of the same, single cell starting form exactly the same initial condition (red dot). This realizations perspective corresponds to the Langevin equation description. The right histogram represents an approximation of the corresponding distribution over the realizations. (b) The picture represents the time evolution of a hypothetical probability distribution. A population of cells initially presents a narrow distribution centered at an intermidiate state value: most cells have an intermidiate state and no individuals show an extreme (low or high) value. As time evolves the shape of the distribution changes – gets wider – , and the population reaches lower and higher values. This perspective corresponds to the Fokker-Planck equation description. (c) A cell can follow different paths (grey dotted lines) to reach a final state *x_f_* starting from an initial state *x*_0_. A finer quantitative characterization of the specific transition from state *x*_0_ to state *x_f_* in terms of highly probable paths and difficulty of differentiation processes can be gained by means of calculating a dominant path (red line) for the transition using a path-integral formalism. For simplicity, the cell state is represented by one variable *x* in all three cases.

To summarize this section: when a stochastic component with specific properties is introduced in a continuous-time dynamical model of developmental dynamics the behavior of the system can be studied from different, mathematically equivalent perspectives; one of the perspectives could be more appropriate than the others given the biological question of interest; the different perspectives complement each other, nonetheless. It is important to note that the three approaches mentioned above (e.g. Langevin, FPE and path-integral) although just recently introduced in systems biology [53,54,80]; are actually well established tools in nonequilibrium statistical mechanics and the stochastic approach to complex systems [71,76,81].

### A Diffusion Approach to Study the EL

The three perspectives to study continuous-time stochastic models of developmental dynamics, briefly described in the previous subsection and represented in Figure 4, have been applied to the understanding of actual developmental cases from an EL point of view. Villarreal and collaborators recently proposed a procedure to construct a probabilistic EL by calculating the probability distribution of stable gene expression configurations arising from the topology of a general N-node GRN [53]. In this approach, the focus of study is the temporal evolution of the distribution over state space (Figure 4b) starting from a position centered on a specific attractor configuration. In other words, the proposed framework predicts how a cloud of cells distributed over a particular attractor will diffuse in time to the neighboring regions (attractors) in state space, given a specific GRN (which constraints the state trajectories). The method has been applied to the case of early flower morphogenesis (see subsection above); and its behavior, in both wild-type and mutant conditions, was in agreement with the temporal developmental pattern of floral organs attainment in *A. thaliana* and most flowering species [53]. This implementation, although consistent with the conceptual basis of the EL, does not depend on an underlying (quasi)-potential function and uncovers the interplay of nonlinear constraints and stochastic forces driving developmental dynamics.

The existence of a potential or “potential-like” function associated with diffusive systems, on the other hand, has been an intensive focus of study in theoretical physics and applied mathematics. Ao and co-workers have published important contributions on this regard [82–85]. Particularly, Ao has proposed a transformation that allows the definition of a function *U*(x) which succesfully acquires the dynamical meaning of a potential function. The corresponding approach has been applied successfully to study several biological systems such as the *phage lambda life cycle* [86], and the carcinogenesis processes, [87,88] from a landscape perspective. This transformation has also been discussed recently in the context of general methods for the decomposition of multivariate continuous mappings *F*(x) and their associated quasi-potentials [28]. These, as well as some associated problems, are still an open area of theoretical research [28,89]. From the available decomposition methods, the one that has been applied the most to specific developmental processes is the potential landscape and flux framework proposed by Wang and co-workers [90]. In this framework, the continuous dynamical mapping *F*(x) is decomposed into a gradient part and a flux, curl part (for details, see [91]). This approach has been applied, for example, to the study of the yeast cell cycle [69,92]; a circadian oscillator [93]; the bi-potent stem cell differentiation process [54]; and neural differentiation [52]. More recently, this method has been applied in the context of the differentiation and reprogramming of a human stem cell network [23]. Here we further discuss the latter as a diffusion landscape approach to study stem cell differentiation.

### Cell Fate Decisions in the Human Stem Cell Landscape

Recently, Li and Wang adopted the diffusion approach to study a previously published [94] human stem cell developmental network composed of 52 genes [23]. In this study they showed how the three perspectives represented in Figure 4 can complement each other in the study of cellular differentiation: (1) through the numerical analysis of the Langevin-like equations for the complete network they acquired a landscape directly from the statistics of the trajectories of the system; (2) by means of approximations they studied the evolution of the probabilistic distribution and obtained an steady-state distribution; and (3) they calculated the dominant paths following a path-integral formalism [80]; the obtained paths were interpreted as the biological paths for differentiation and reprogramming [23]. As Li and Wang showed, from the results of the three perspectives it is possible to quantitatively describe the underlying landscape. One then may be interested in how the landscape changes in response to specific perturbations. A general problem of concern in stem cell research is why do the known reprogramming strategies, which commonly consist on combining perturbations to specific transcription factors, actually work. Li and Wang systematically tested which genes and regulatory interactions affect the most the quantitative properties of the landscape (*e.g*. height values and transition rates) when perturbed. Interestingly, several biological observations associated with the manipulation of the so-called Yamanaka factors (Oct3/4, Sox2, Klf4, c-Myc) - the transcription factors considered the core regulators in the induction of pluripotency - were consistent with the observed modeling results; for example, simulated knockdown perturbations to these factors consistently increased (lowered) the probability (height) of the differentiation state. On the other hand, the path-integral formalism allowed them to show how specific perturbations to these factors cause the differentiation process to be easier or harder in terms of the time spent during transitions and the characteristics of the differentiation paths. In other words, this study presented an important contribution towards the mechanistic, dynamical explanation of the characterized reprogramming strategies in terms of the properties of the underlying EL.

## Concluding Remarks

For the practical implementation of EL models W. Wang recently proposed the following overall four-steps strategy [72]: (1) establishment of a GRN; (2) characterization of the attractor (and quasi-potential) landscape through dynamical modeling; (3) computational prediction of cell state responses to specific perturbations; (4) analysis of the prevailing paths of cell fate change. The first step (1) is already a well-established research problem that includes expert curation of experimental data and/or statistical inference. In this review we focused on the second step and presented examples of how steps (3) and (4) can be achieved once a EL models is effectively constructed. As shown here, there are several ways to implement an EL model starting from a GRN. The specific choice should be made considering the properties of the network and the associated questions of interest. The methodologies reviewed here are mostly well-suited to approach the problem of differentiation at the cellular level in a mechanistic setting. The observed behavior results from constraints given by the joint effect of nonlinear regulatory interactions and the inherent stochasticity; however, the actual physical implementation of these generic mechanisms in a multicellular system would imply additional sources of constraint. Tissue-level patterning mechanisms such as cell-cell interactions; chemical signaling; cellular growth, proliferation, and senescence; inevitably impose physical limitations in terms of mechanical forces which in turn affect cellular behavior. This would thus imply non-homogenous GRNs with contrasting additional chemical and physical constraints. Given this fact, the next logical step to extend EL and associated dynamical models would be to account for these physical processes in an attempt to understand how cellular decisions occur during tissue patterning and not just in cell cultures. Although some progress has been presented in this direction [95,96], the problem certainly remains open. From a theoretical perspective, a further challenge would be to carefully evaluate the assumptions implicit in the EL models. For example, the adoption of the diffusive perspective briefly explained above - which is often taken as a standard in stem cell systems biology - implicitly assumes certain properties about the forces driving the temporal evolution of the system [81]. Do these conditions are universally met by developmental systems? Recent interesting work is starting to suggest the biological relevance of additional constraints such as state-dependent fluctuations [97,98].

Overall, the application of the methodologies discussed in this review to specific developmental processes has shown the practical relevance of dynamical models consistent with the conceptual basis of the EL and the fundamental role of the constraints imposed by the GRN interactions. Both approaches, landscape-(in)dependent, are useful to answer specific questions and can complement each other. So far, EL models have shown to be an adequate framework to understanding stem cell differentiation and reprogramming events in mechanistic terms; and are also starting to show promise as the basis for rational cancer therapeutic strategies.

## Acknowledgments

This work was supported by grants from CONACYT, Mexico: 124909 to ERAB; from PAPIIT-UNAM, IN229009-3 (ERAB). JDV receives a Phd scholarship from CONACYT. The authors acknowledge technical support of Rigoberto V. PerezRuiz and logistical and administrative help of Diana Romo. The authors aknowledge the Centro de Ciencias de la Complejidad (C3), UNAM.

